# Biocontainment of phages inhibits bacterial clearance in micro niches

**DOI:** 10.64898/2026.07.02.736089

**Authors:** Liam Boot-Handford, Remy Chait, Tobias Bergmiller, Herve Migaud, Charles R. Tyler, Ben Temperton

## Abstract

Phage therapy offers a promising solution to the antimicrobial resistance crisis. However, a major concern preventing the adoption of phage therapy is the potential for unintended consequences of phage release; both in regard to preventing the spread of phage resistance, and the proliferation of a non-endemic virus into the microbial ecosystem. Conditional replication (biocontainment) of phages through bioengineering may address these concerns, but the impact on bactericidal efficacy is unknown. Here, we created a biocontained T7 phage (T7Δcapsid) lacking the major structural capsid gene, gp10AB, that can only replicate on Escherichia coli strains expressing gp10AB in trans, and assessed its bactericidal efficacy compared with wild-type T7. Congruent with model predictions, T7Δcapsid was only able to clear a well-mixed culture of E. coli at a multiplicity of infection (MOI) of 10 or higher, whereas wild-type T7 prohibited growth at an MOI of 0.1. The reduction in efficacy was more evident in a complex structured environment within a microfluidic device, where phage success depends on its ability to penetrate a microbial niche via propagation. In this environment, T7Δcapsid was unable to propagate into the bacterial population and unlike wild-type T7, had no impact on the population’s growth. This study shows that whilst biocontainment of phages may improve the biosafety of phage therapy, it comes at the cost of its propagation efficacy and niche penetration in relevant environments.

## Introduction

Antimicrobial resistance (AMR) is an escalating global issue which greatly threatens human health and modern medicine, and novel or innovative antimicrobial therapeutics are urgently needed. Bacteriophages (phages) are natural viruses which infect and kill bacteria regardless of their AMR status. Phages are the most abundant biological entity on earth at 10^31^ individual virions^1^, and they have been used safely as adjuncts or alternatives to pharmaceutical antibiotics^2,3^ since their first characterisation in 1915 (4). Their functionally limitless diversity, unique ability to replicate at the site of infection, high potential for engineering and synthetic biology^5–7^, and potential as precision medicines give them numerous, indispensable advantages in the fight against drug-resistant bacteria.

A potential concern of phage-based therapies is their potential for escape and proliferation in the natural environment, during or after their use. This is founded on two legitimate biosafety concerns: (1) injudicious use of phages could repeat historical mistakes of small-molecule antibiotics, where consistent environmental exposure to antimicrobials promoted resistance in bacteria, both in clinical and environmental applications^8,9^; (2) potential unintended consequences of releasing non-endemic species into an ecosystem where invasive species could threaten natural diversity^10,11^. These concerns are heightened when phages are engineered to be more effective as therapeutics, due to the potential of increased fitness^12,13^ and subsequent potential disruption of the microbiome of the natural environment and biota^14^. Deliberate release of genetically modified organisms is strictly regulated in the European Union^15^ and United Kingdom^16^, requiring stringent compliance with broad criteria such as traceability. Due to the nanoscopic size and self-replicating nature of phages, controlling and tracking their proliferation via filtration or chemical treatment is challenging.

Biological containment (biocontainment) refers to measures taken to prevent replication of an organism outside of a controlled environment, and to ensure there is no exchange of transgenic DNA^17^. Secure biocontainment could provide the reassurance that regulatory bodies need to approve phages as therapeutic or prophylactic agents in clinical, agricultural (e.g. crop spraying^18^) or aquaculture settings (e.g. immersion delivery directly into water^19^). Previously, biocontainment of phages has been achieved through two main methods of bioengineering: deletion of essential phage genes encoding a critical structural protein such as the capsid^20^; or recoding start codons within structural genes to amber stop codons (AUG to UAG)^21^. These engineered phages can only produce viable phage progeny and replicate on specific bacterial host strains that either complement the essential phage protein *in trans*^20^; or express amber-initiator tRNA and matching tRNA synthetases to decode amber codons to start codons^21^. Such biocontained phages can still inject their packaged DNA and execute their lifecycle to kill host cells which lack these accessory factors, but no viable phage progeny are produced from these infections. Thus, biocontained phages can only sustain a single round of infection outside of specific propagation host strains, alleviating concerns of environmental escape, as any biocontained phages would either be sequestered by infection of the target or would degrade over time^22^. Removal of auto-catalytic, self-propagating features enables biocontained phage therapy to be more comparable to chemical compounds in regard to dosing, environmental impacts and escape, whilst maintaining the unique advantage of host specificity.

A major caveat to the use of biocontained phages is that it removes one of the most powerful and unique features of phages as antimicrobials – their ability to “auto-dose” via self-replication at the site of infection^23^. In an idealised scenario, as long as one viable phage particle reaches a target pathogen, the localised production of tens to thousands of new phage particles^24^ per round of infection creates a chain reaction that eradicates bacteria, and self-limits once no more prey items are available. This chain reaction causes a propagation wave that carries the infection beyond its initiation as neighbouring cells are infected and destroyed – a feature which is demonstrated in plaque and spot assays^25,26^, and in motile bacteria^27^.

Overcoming this limitation requires increasing dosing of biocontained phage using high phage-to-host cell ratios (also referred to as multiplicity of infection, or MOI) to ensure bacteria are eradicated in a singular infection wave. In scenarios where the bacteria are distributed within a well-mixed system and phages can be quickly and evenly dispersed, this is feasible: Biocontained phages have been shown to be as effective as WT phages to treat sepsis in mice at an MOI of ∼500^20^. Similar efficiencies would also likely scale to using phages for clearance of planktonic bacteria within recirculating aquaculture systems.

However, in cases where the bacterial hosts are embedded within highly structured environments or biofilms, such as the gut, the mucosal microbiome, or wounds, this approach may be limited. In such environments bacterial populations are highly stratified by physical and physiological constraints, and protection of bacterial populations via biofilm formation is a common cause of resistance to small-molecule antibiotics^28^. It is estimated that due to the fitness benefits, 80% of all bacteria and archaea on Earth exist in biofilms/structured assemblies rather than as planktonic cells^29^. Replicating phages can still penetrate multi-layered communities via repeated infection cycles that cascade from the exposed cell-layer^30^, with bacterial clearance a function of the number of exposed cells, phage infection dynamics and bacterial growth kinetics^31–33^. If the bacterial growth rate and population size exceeds a clearance threshold determined by the phage infection kinetics, the bacterial population will persist^34^. Therefore, in these ‘micro niches’ where the phage success is dependent on ability to penetrate into a shielded community such as in a biofilm^30,35^, or in the mucosal microbiome^36^, biocontained phage therapies would face an inherent limitation which cannot be mitigated by escalating the dose. In these cases, replicative phages may be more effective due to their ability to concentrate around the target site^31^.

Here, we evaluated the trade-off between biocontainment and niche-penetration by engineering a biocontained T7 phage (T7Δcapsid) via deletion of genes encoding the major and minor capsid proteins, *gp10AB*, that can only propagate on a bacterial host expressing gp10AB from a plasmid *in trans* (Figure 1, panels i-iii). We first mathematically modelled bacterial clearance at MOI 0.1, 1, 10 and 100 using biocontained phages in well-mixed experimental cultures compared to replicative phage. This confirmed the assumption that an MOI of ≤1 would be insufficient to clear a replicating bacterial population for T7Δcapsid. MOI of ≥10 (where Poisson distributions predict that >99% of cells are expected to have at least 1 phage adsorbed) resulted in complete bacterial clearance. Model predictions were compared to *in vivo* experiments run at the same MOI, to confirm model accuracy. We next evaluated the ability of T7Δcapsid to penetrate a multi-layer bacterial niche on a custom microfluidics device (Chait, communication) to assess penetration and clearance of micro niche environments, where phage exposure was limited to a few cells atop a microbial population (Figure 1, panel iv). In this environment, phages were unable to migrate into the niche, preventing clearance. Our findings suggest that while biocontainment of phages is an attractive proposition to address regulatory concerns around their environmental escape, the potential associated reduction in efficacy may severely limit their application for disease management outside of well-mixed bacterial populations.

**Figure 1:**
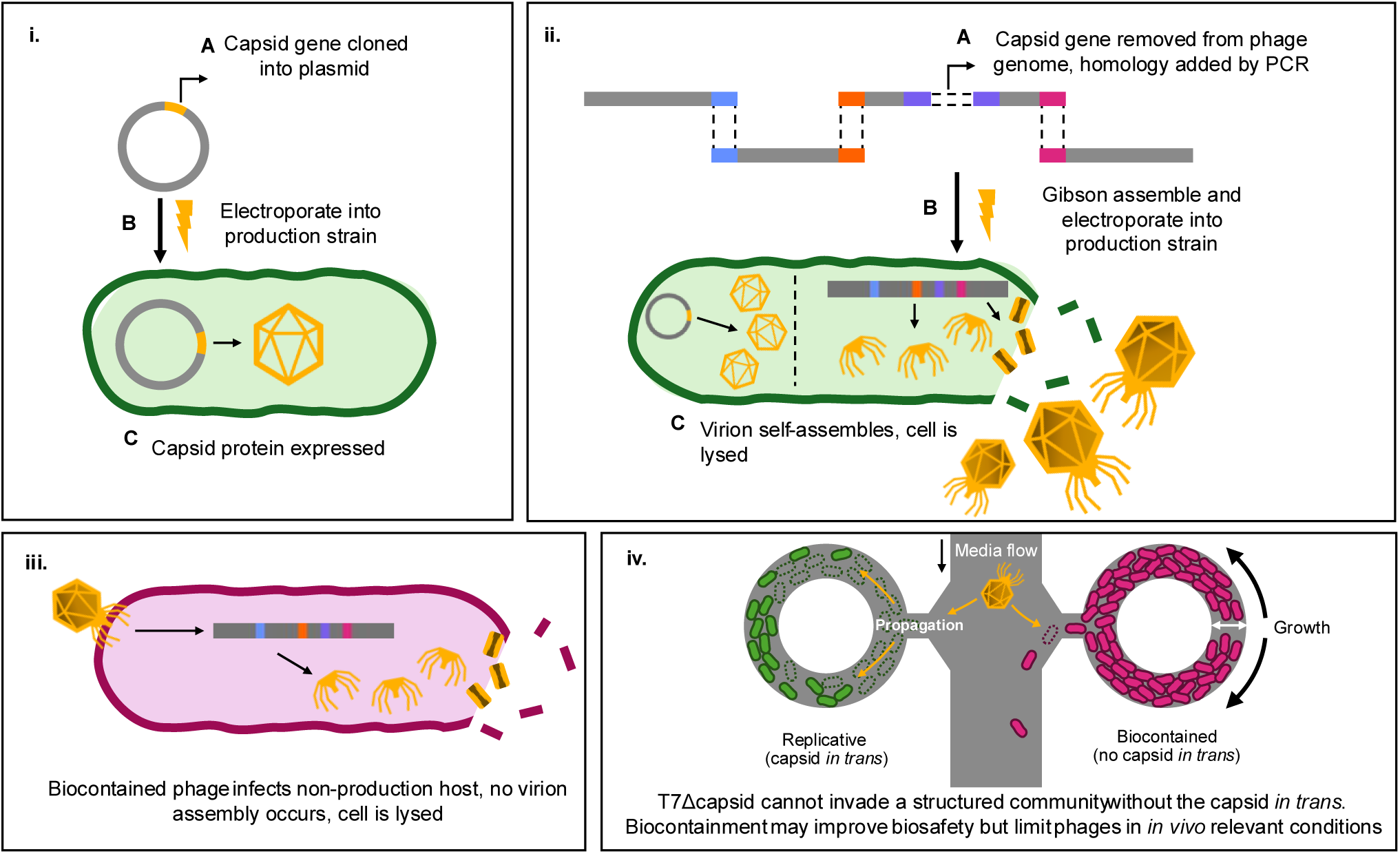
biocontained T7 is made via *in trans* gene complementation, genome Gibson Assembly, and evaluated in a custom microfluidic device. (i) The capsid gene *gp10* is cloned into a plasmid, and electroporated into the production strain. (ii) The phage genome is amplified via PCR, omitting *gp10* and including 40 bp orthogonal overlapping homology. *E. coli* harbouring the *gp10* plasmid is transformed with Gibson assembled T7Δcapsid genome, and virion assembly proceeds. (iii) When infecting a non-production host, T7Δcapsid cannot assemble an entire virion and the cell is lysed without releasing progeny. (iv) The ability of T7Δcapsid to invade a bacterial niche is tested in a custom microfluidic device. Bacterial killing is compared between replicative (capsid *in trans*) and biocontained (no capsid *in trans*) strains within the same conditions.

## Materials and methods

We produced T7Δcapsid via Gibson assembly in order to test its antibacterial efficacy in well mixed liquid culture, and an *in vivo*-relevant model of a bacterial niche (Figure 1, panel iv). This was compared to the mathematical prediction of bacteria and phage dynamics when T7 is biocontained or replicative. If accurate, this may be able to predict the effect of biocontainment on the efficacy of other phages, which could prevent the unnecessary engineering of inefficient phages.

### Mathematical modelling of infection in well-mixed cultures

Biocontainment of phages can be modelled using established ordinary differential equations^37^ by setting the term for burst size (*β*) to 0. The rate at which bacterial cells become infected and release new viral progeny to propagate the infection can be modelled as follows:

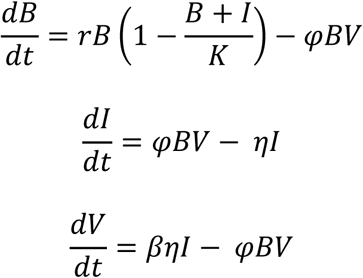

where B is the number of uninfected cells, r is the maximum growth rate of the bacteria, I is the number of infected cells, *φ* is the adsorption rate, V is the number of free viruses, β is the burst size, η is the lysis rate and the mean infection period is 1/η.

These equations were used with the *solve_ivp* function within the scipy python package to evaluate phage and bacterial dynamics in well-mixed, nutrient replete cultures at various MOIs (0, 0.1, 1, 10, 100), using the following parameters:

### Phage genome and plasmid preparation

A plasmid was assembled via Golden Gate assembly combining the p15A origin and ampicillin resistance gene with *gp10AB* (p15A_Capsid)^38^ (Supplementary Table S1. Commercial chemically competent cells of *E. coli* strain DH10β (New England Biolabs) were transformed with this plasmid. A control plasmid was also created with the same origin and resistance marker, but without gp10AB (p15A_Control). Both sequences were confirmed by nanopore sequencing and assembly (PlasmidsNG, UK) and transformed into two *E. coli* strains: TB204 (Addgene #230034) and TB205 (Addgene #230033), which contain sfGFP and mCherry fluorescent genes respectively, under constitutive PL promotors (Supplementary Table S2).

T7Δcapsid was constructed by amplification of five fragments via PCR, with 40bp homologous overlaps, omitting *gp10AB* (Supplementary Figure S1, Table S3). Gibson assembly (New England Biolabs) was used to assemble this into a full-length phage genome according to the manufacturer’s instructions. The assembly was diluted to 100 µL in nuclease free water (NFW) and mixed with 100 µL 100% isopropanol, incubated at room temperature for 20 minutes, and centrifuged at 15,000 × *g* for 20 minutes. The supernatant was discarded and the pellet washed in 200 µL 70% ethanol, centrifuged again, and the pellet was allowed to dry before resuspension in 5 µL NFW for transformation.

### Transformation

Chemically competent *E. coli* TB204 harbouring p15A_Capsid was prepared for phage genome transformation following an adapted protocol^39^: A culture of TB204/p15A_Capsid was grown from a single colony in LB with 100 µg/mL ampicillin, 10 mM MgCl_2_ and 10 mM CaCl_2_ (LB_amp_) (37 °C, 200 rpm) until stationary phase, then 500 µL of this culture was inoculated into 25 mL of LB_amp_ and grown for 6-10 hours at 22 °C, 200 rpm. Once the culture reached an OD_600_ of 0.4-0.5, it was chilled on ice and centrifuged at 4000 × *g* at 4 °C for 15 minutes. The supernatant was discarded, and the pellet washed twice in 12.5 mL of ice cold 0.1M CaCl_2_, before being resuspended in 200 µL ice cold 0.1M CaCl_2_. An aliquot of 50 µL of chemically competent cells was co-incubated on ice with 5 µL of assembled phage genome for 30 minutes, heat shocked at 42 °C for 30 seconds and placed back on ice. After 5 minutes, 950 µL of SOC (New England Biolabs) was added at room temperature and immediately mixed with 3 mL top agar (0.65% w/v bacteriological agar, 10 mM MgCl_2_, 10 mM CaCl_2_, 100 µg/mL ampicillin). This bacterial top agar suspension was poured immediately onto a pre-warmed agar plate. Once dried, the agar plate was incubated at 37 °C overnight. Plaques were picked and purified using a previously published method^40^. Briefly, this involved removing an agar core of a single plaque, propagating the suspended phage on *E. coli* TB204/p15A_Capsid, then diluting the lysate to single plaques and repeating the process twice more. The final phage core was propagated on *E. coli* TB204/p15A_Capsid and the lysate filtered through 0.22 µm and stored at 4°C. The genome sequence of T7Δcapsid was confirmed by short read sequencing (MicrobesNG, Illumina 2 × 150 bp) and assembled as described previously^40^.

### Spot and liquid culture growth assay

Spot assays were performed to confirm biocontainment of T7Δcapsid. Purified phage lysate from T7Δcapsid propagated on TB204/p15A_Capsid was spotted onto top agar bacterial lawns of either TB204/p15A_Control, or TB204/p15A_Capsid (6 mL top agar, 2 mL bacterial culture at OD_600_ = 0.6), in 5 µL droplets at serial dilutions down to 1 × 10^−11^, with plates incubated overnight at 37 °C.

Phage virulence in liquid culture was assayed in 96 well plates containing a mixture of TB204/p15A_Capsid or TB204/p15A_Control cultures diluted 1/100 from OD_600_ of 0.55 and T7Δcapsid at MOI from 100 to 0.1. Plates were incubated in a TECAN Sunrise plate reader at 37 °C for 24 hours and the OD_600_ measured every 15 minutes. OD_600_ for all treatment wells was normalised against LB_amp_ blank media within the same plate, and positive controls of both bacterial strains were included with no phage.

### Micro-niche assay

Microfluidic devices were used to determine T7Δcapsid efficacy in invading host populations in bacteria-scale physical microniches (Figure 1, panel iv). Briefly, toroidal growth chambers (channel width approximately 6 µm, height approximately 1 µm), are arranged alongside and connected to a larger flow channel from which they are fed by diffusion of nutrients and into which excess cells from the growing population are washed away. We have found these devices are robust in operation over long timescales and lack fitness-determining features such as corners where “founder” cells can get trapped. The chambers and main flow channel were formed by soft replica moulding of polydimethylsiloxane (PDMS, Corning Sylgard 184, 1:10 ratio) against SU-8 epoxy photoresist features produced by photolithography on a silicon wafer. The PDMS was cured at 70°C overnight, inlets punched using sharpened flat-faced cannulae and then bonded by air plasma to isopropanol-cleaned dry glass coverslips.

In use, the growth chambers were seeded by flooding the device with liquid overnight cultures of *E.coli* TB204/p15A_Capsid and TB205/p15A_Control mixed in 1:1 ratio and suspended in LB, containing 100ug/ml ampicillin and 0.2% v/v TWEEN-20. Cells flowed into the growth chambers with the media. Filter sterilised LB containing 100µg/mL ampicillin + 0.02% v/v TWEEN-20 was then pumped through the device at a rate of 4 mL/hour for 30 minutes to clear cells from the main channel and tubing. The flow was then reduced to 1.5 mL/hour and the device incubated at 30°C for 24 hours. Once single strain well populations were established, LB containing 100µg/mL ampicillin, 0.02% v/v TWEEN-20, and 4.8 × 10^6^ T7Δcapsid PFU/mL was pumped through the device at a rate of 1.5 mL/hour for 20 hours. T7 WT was then mixed with LB containing 100 µg/mL ampicillin + 0.02% v/v TWEEN-20 and pumped through the device at a concentration of 4.8 × 10^6^ PFU/mL for 7 hours. GFP (Ex:509nm, Em:) and mCherry (Ex:610 nm Em:) fluorescence and phase contrast images were acquired every 5 minutes across 10 chamber pair locations. Growth chambers were selected where a single strain was established in each chamber. The images were analysed and timelapse videos created using ImageJ (version 1.54p). Briefly, the fluorescent images were shading-corrected and all images registered across the time series using custom Matlab scripts^41^. Fluorescent images were binarized and intensity within the chamber summed to provides quantitative data on the population occupancy of each chamber. Finally, all fluorescent data were expressed as a fraction of peak intensity for data visualisation, calculated as the peak fluorescence within the pooled mean fluorescence across all chambers.

## Results

### Spot assay confirming biocontainment of T7Δcapsid

T7Δcapsid was created to use as a model to assess the efficacy of biocontained phages in killing bacteria. T7 was chosen as a model phage for this assay as it is one of the most characterised phages^42^ and can be biocontained by deletion of the capsid gene *gp10AB* and propagated on strains containing a plasmid encoding *gp10AB* downstream of a T7 native promoter^20^. The genome of T7Δcapsid does not contain the capsid gene, *gp10AB*, and can only replicate within a host which expresses *gp10AB* from a plasmid. To confirm biocontainment of T7Δcapsid, zones of lysis (ZOL) were compared between T7Δcapsid and T7 WT, when infecting TB204/p15A_Capsid and TB204/p15A_Control (Figure 2). T7Δcapsid reached 1.46 × 10^8^ PFU/mL when propagated on TB204/p15A_Capsid, which constitutes an approximate ten-fold reduction compared to WT propagated on the same strain. There was also no visible ZOL at dilutions of 10^−3^ or greater on TB204/p15A_control. Under restricted replication on TB204/p15A_control, ZOL formed by T7Δcapsid lacked a ‘halo’ morphology usually associated with wildtype T7.

**Figure 2:**
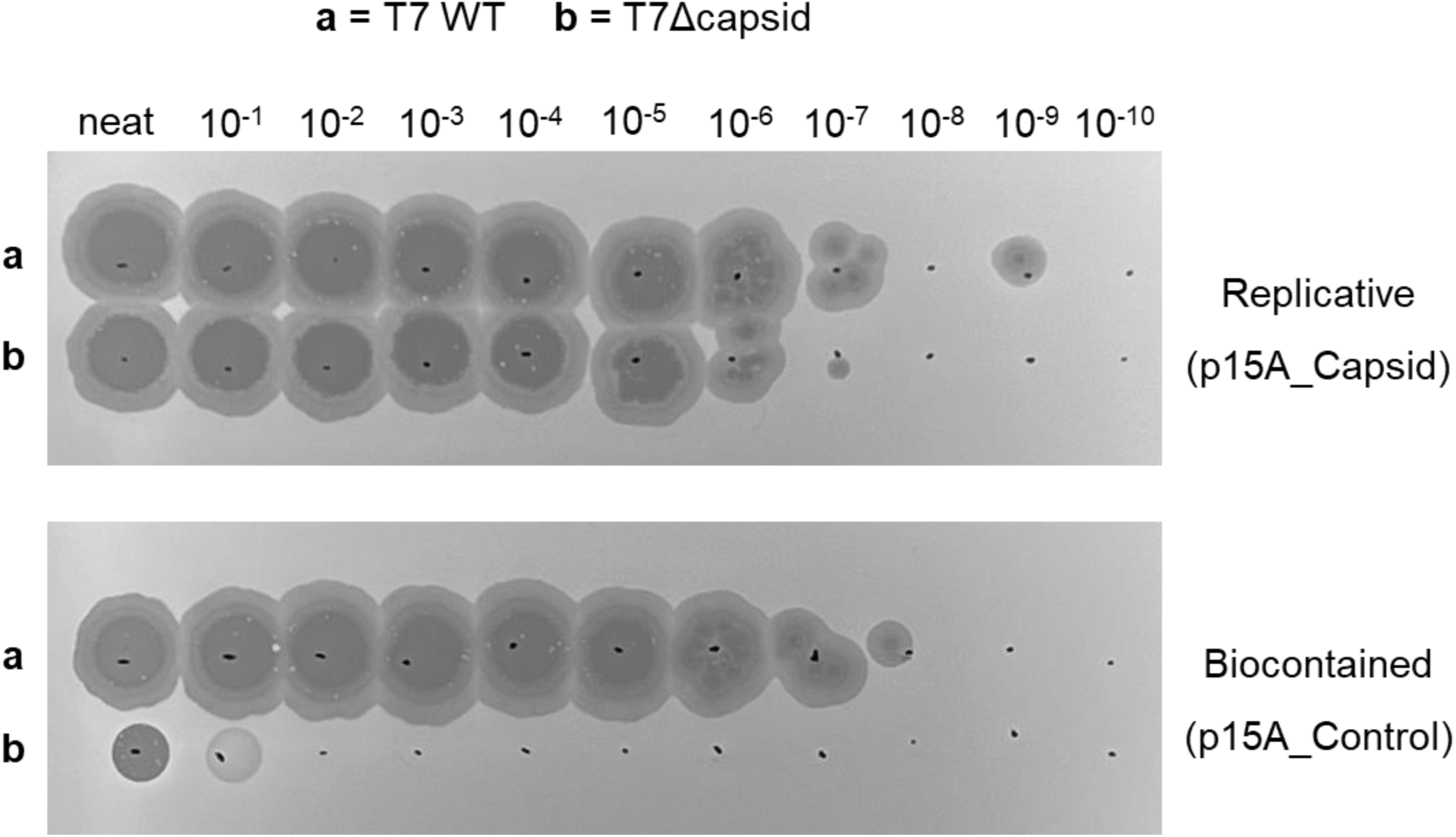
biocontained T7 has a modified zone of lysis morphology. Phage stocks of T7Δcapsid and T7 wild type (WT) are spotted onto a lawn of replicative (p15A_Capsid) or biocontained (p15A_Control) host.

### Mathematical model of T7Δcapsid infection correlates with empirical data

A previously published mathematical model was used to predict the effect of biocontainment (burst size = 0) on T7, when in a well-mixed bacterial suspension at increasing MOI^37^. The equations were used with the *solve_ivp* function within the scipy python library using the parameters available in Table 1. This model predicted that at an MOI of ≤ 1, the bacterial population would survive exposure to phage (Figure 3, inset). In the prediction of replicative phage, eradication of bacteria was achieved at MOI ≥ 0.1.

**Figure 3:**
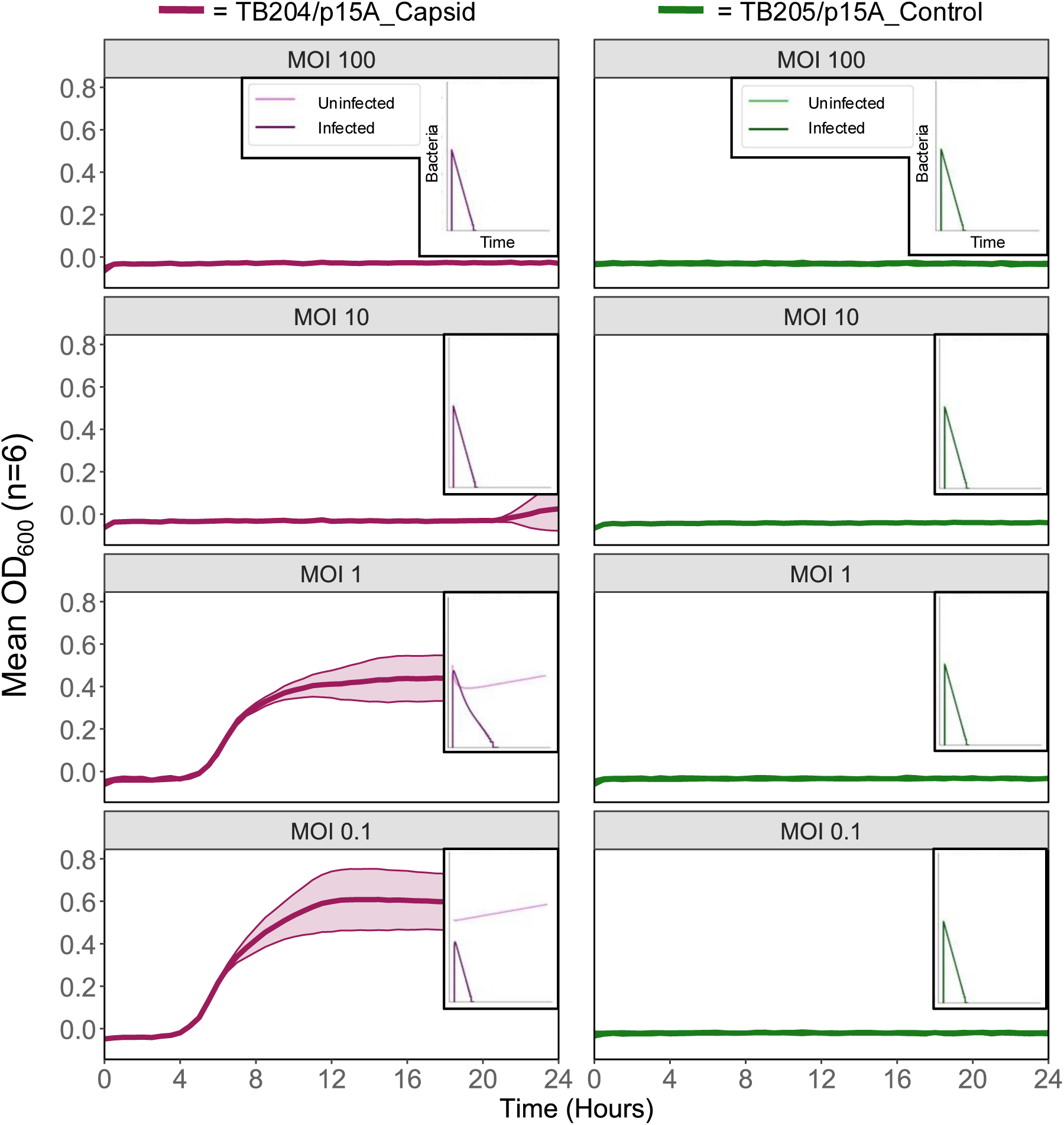
MOI ≤ 1 is required for biocontained phage to eradicate a well-mixed population. **Figure inset:** Mathematical modelling of phage and bacterial interactions in a well-mixed population where phages are either wildtype (burst size = 200) or biocontained (burst size = 0). **Figure main:** Absorbance at OD = 600nm for E. coli TB204/pControl (left) and TB204/pCapsid (right) when mixed with T7Δcapsid at decreasing MOI and incubated in LB_amp_ for 24 hours at 37 °C.

**Table 1:**
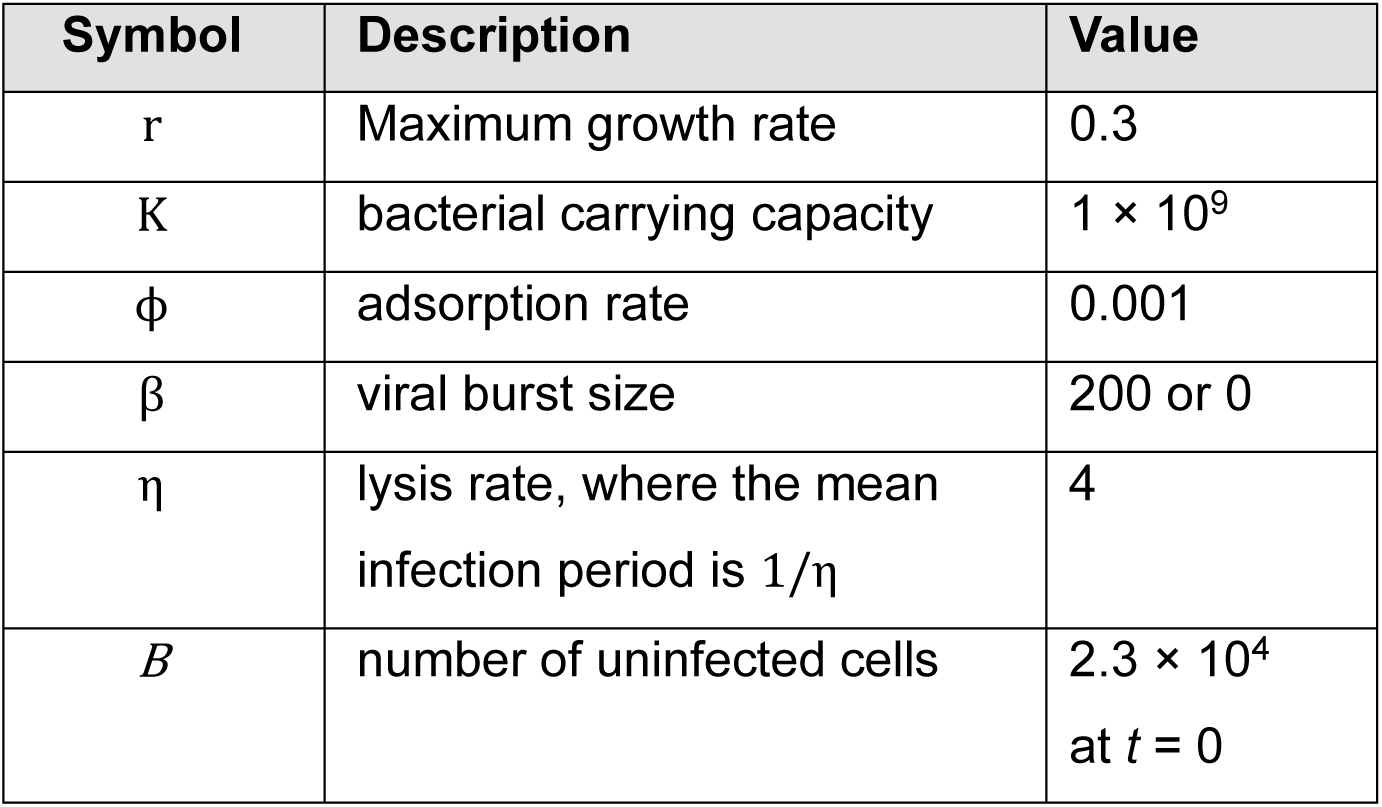
parameters used to simulate biocontained and replicative phage.

Next, we set up bacterial liquid growth assays using the same MOI range as in the mathematical model (10^−1^ – 10^3^) and measured the bacterial growth using OD_600_ for 24 hours. This showed that the mathematical model could predict the outcome of exposure to T7Δcapsid. T7Δcapsid was unable to replicate after infecting TB204/p15A_Control, and as a result it was unable to eradicate populations of bacteria at MOI ≤ 1 (Figure 4). Where replication was restored within TB204/p15A_Capsid, T7Δcapsid prevented growth of bacteria at all MOI for 24 hours. After approximately 22 hours of exposure to T7Δcapsid, growth began at MOI = 10 in one of six replicates, potentially due to emergent phage resistance.

**Figure 4:**
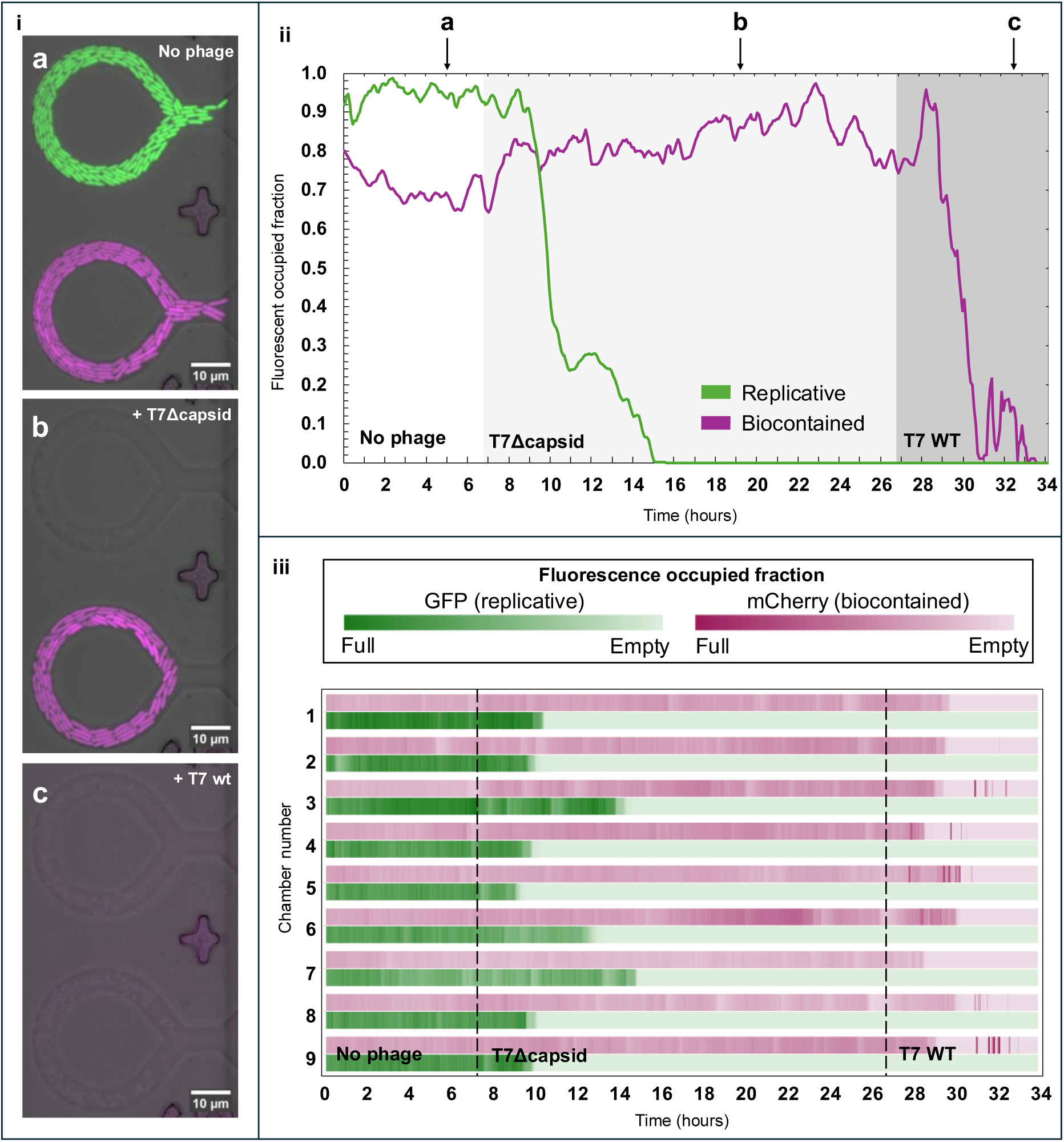
Biocontained phage T7Δcapsid cannot invade a bacterial microniche without replication enabled by capsid complementation. Differentially fluorescently-labelled *E.coli* hosts, grown side-by-side in microfluidic niches, either constitutively express T7 capsid (TB204(GFP)/p15A_Capsid, green), enabling replication of capsid-deleted T7 phages, or contain empty expression vector (TB205(mCherry)/p15A_Control, magenta) that maintains biocontainment of the phages. After phage-free growth (i, a), exposure to phage T7Δcapsid enables invasion and clearance of replication-permissive but not biocontained host niches (i, b). T7Δcapsid-biocontained niches are subsequently invaded and cleared by non-biocontained wild-type phage T7 (i, c). Fluorescent cell occupancy (median smoothed fraction) of micro-niches during growth and phage exposure (ii) shows invasion and rapid clearance only when invading phages can replicate. Fluorescent cell occupancy arranged by adjacent niche pairs, shows variation in time between phage exposure and bacterial clearance (iii).

### T7Δcapsid cannot invade a bacterial micro niche

The ability of T7Δcapsid to penetrate a bacterial micro niche was assessed in microfluidic growth chambers, where only the outermost layer of the bacterial population is exposed to the phage. Survival of bacterial populations in monoculture growth chambers was measured for an equal number of chambers containing TB204/p15A_Capsid or TB205/p15A_Control, when exposed to T7Δcapsid.

T7Δcapsid was unable to invade bacterial micro niches when biocontained. Exposing growth chambers populated with TB205/p15A_Control to T7Δcapsid showed no measurable reduction in growth after 16 hours of exposure (Figure 4). In contrast, growth chambers containing TB204/p15A_Capsid were eradicated within 8 hours of exposure. The time between initial exposure to T7Δcapsid and eradication varied between 2 and 8 hours across all chambers (see Supplementary video 1). There was bright fluorescence signal in the mCherry channel after all TB205/p15A_Control cells had been killed, however this did not correlate to cellular growth (Figure 4, panel iii) and was potentially caused by fragments of biofilm or cellular debris traveling through the main channel. This was not observed in the sfGFP expressing strain. A tenth growth chamber, not included in Figure 4, had a mixed population of both strains, and in this case T7Δcapsid was sufficient to clear the chamber of bacterial cells (Supplementary video 2).

## Discussion

Biocontainment of phages may reduce barriers to regulatory acceptance as novel antimicrobials by reducing the risks of environmental escape and accumulation. However, auto-dosing of phages is a unique and important feature of phage efficacy. As predicted by mathematical models^37^, T7Δcapsid empirically had a reduced antibacterial effect compared to wildtype T7 in all models tested. We only observed bacterial eradication when MOI was >10 in both well-mixed cultures and via spot assays. In well-mixed co-cultures of bacteria and phage, all cells are equally likely to encounter a phage particle, and thus bacterial killing is a function of cell and viral density. When MOI > 10, there is a 99.5% probability that each cell encounters at least one phage, after which normal phage infection and lysis occur with both wildtype and biocontained T7. Wildtype T7 has a burst size of ∼200 progeny per infection^43^, and so even at very low starting MOI, viral concentration increases rapidly after only a few rounds of replication. In biocontained phages, the effective burst size is zero as no viable phage progeny are produced.

Well-mixed cultures are a poor proxy for many clinically relevant bacterial infections, which often form biofilms and/or inhabit micro niches^44,45^. To better capture these dynamics, we designed toroidal microfluidic growth chambers containing bacterial populations with limited environmental exposure. Within these growth chambers, biocontained T7Δcapsid did not reduce the bacterial population of non-permissive TB205/p15A_Control cells. In contrast, all TB204/p15A_Capsid were eradicated within 8 hours of exposure. When the wildtype T7 was added, all remaining cells – which were exclusively TB205/p15A_Control – were killed within 5 hours. Previous work has shown that microfluidic devices can be used to model phage-susceptible and resistant populations of bacteria^32^, which was not seen here – likely due to the smaller population sizes within the growth chambers used. Microniche invasion of biocontained phage has not previously been reported.

The onset of population collapse of both TB204 and TB205 strains varied between growth chambers despite simultaneous exposure to phage within the microfluidic device. This is consistent with the expectation that with wildtype phages, propagation can only occur if the infected cell remains within the population of uninfected cells until the release of new viral progeny. The design of the toroidal channels enables cells proximal to the central channel, where phage infection is most likely to occur, to be pushed out by growth from below. The variable time to lysis observed in this study is likely caused by the stochastic interaction between phage encounter rates and bacterial cell wash-out. However, when a cell within the well is lysed, the localised release of progeny enables propagation throughout the population.

Translating the findings of this research into *in vivo* application suggests that the efficacy of using biocontained phage to clear established bacterial infections may be limited. Hypothetically, if a patient was infected by 10^5^ bacteria within the bloodstream, 10^7^ phages would need to be administered to clear the infection at MOI = 100 to eradicate the bacteria. This approach was effective in rescuing mice from the same bacterial load, at MOI = 500^20^. However, the success of this approach may depend entirely on whether the bacteria are embedded within a biofilm - which will then prevent eradication and re-seed the infection upon phage clearance. Efficacy will be further limited by clearance of biocontained phage by the host immune system, as the number of active phage in the blood will never increase beyond the initial dose^46^ whilst the bacterial population can grow. In contrast, a self-replicating phage can self-dose at the site of infection, allowing penetration into a shielded population. In an aquaculture context – where phage therapy has received significant attention^47,48^ – there are applications were biocontained phage could be suitable. For example, biocontained phages could be added when fish rearing or transfer systems are decontaminated in between production cycles. Use in this context as a water treatment may help reduce bacterial load and prevent contamination, without needing regulatory approval as a medicinal product. The relatively small water volumes would also make high phage concentrations more achievable. In these cases, biocontained phage would still require mechanical disruption of biofilms to reduce the availability of shelter for bacteria.

In addition, there would be high selection pressure on biocontained phages to escape biocontainment via recombination with closely related phages^49^, which may be present within a complex community. An alternative mechanism of escape could utilise structural protein piracy, through which a biocontained phage could create infective virions by enlisting the proteins produced by another phage^50^. There is no data to date on how likely this is to happen with a biocontained phage, but the extremely high number of replication events involved in phage propagation to reach a high titre, coupled to the extreme fitness advantage upon success, means the likelihood of escape of biocontained phages must be empirically demonstrated to be effectively zero before regulatory approval.

Through careful consideration of phage selection to reduce genome homology and bioengineering it may be possible to generate biocontainment that cannot escape, whilst having a limited number of rounds of replication. This has been engineered in bacteria^51^, and installing a replication-permissive plasmid into a cell population with a strict ‘expiry date’ could allow a controlled number of phage propagation cycles. All biocontained phage will either infect the target pathogenic host and terminate replication, or infect the introduced replication-permissive host and release biocontained progeny. After the ‘expiry date’ has passed, all replication-permissive hosts will have expired or been lysed, and therefore the phage and bacteria both go extinct. A mechanism of this type could retain the efficacy of a natural phage whilst addressing concerns over environmental stewardship.

This work suggests that biocontainment of T7 through deletion of structural genes may still allow efficient bacterial clearance in well-mixed systems, whilst presenting secondary benefits for regulation through reduced environmental impacts. However, biocontainment impeded a potent feature of phage biology for treating disease, particularly important in established infections that occupy biofilms and micro niches. The impact of biocontainment should be further assessed in *in vivo* infections, as well as using a broader diversity of phages and mechanisms of containment before biocontained phages are pursued above natural phages. Ultimately, biocontained phages may present a complex solution to the problem of antimicrobial resistance, and regulatory reform to include natural phage therapeutics may ultimately be a more tractable path to translatable phage therapy.

## Data availability

NCBI Accession number for T7Δcapsid is provided in Supplementary Materials. All primers, plasmids and strains used are provided in Supplementary Materials.

## Code availability

The figures in this study were created with R code, available at https://github.com/mrliambh/boot-handford_microniche.

## Acknowledgements

The authors would like to acknowledge the contributions of Dr Michelle Michelsen for assistance writing code.

## Author contributions

LBH created T7Δcapsid and all plasmids used, performed the experiments and wrote the manuscript. RC created the microfluidic devices and performed the growth chamber experiments with LBH. TB created the fluorescent strains. RC, TB, HM, CRT and BT edited the manuscript.

## Competing interests

None.

## Funding

LBH is undertaking a PhD studentship supervised by BT, CRT and HM and funded by Mowi Scotland Ltd.

## Abbreviations

MOI: Multiplicity of infection
AMR: Anti-microbial resistance
NFW: Nuclease-free water

## Notes

### Competing Interest Statement

The authors have declared no competing interest.

