## Supplementary materials for "Biocontainment of phages inhibits bacterial clearance in micro niches"

Table S1: Plasmid details used in creating and assaying T7Δcapsid.

| **Plasmid name** | **Backbone** | **Resistance** | **Promoter** | **Gene** | **Function** |
| --- | --- | --- | --- | --- | --- |
| **p15A_Capsid** | p15A | Ampicillin | T7 native promoter | *gp10AB* | Permits replication of T7Δcapsid |
| **P15A_Control** | p15A | Ampicillin | None | None | Confers amp resistance without permitting replication of T7Δcapsid |

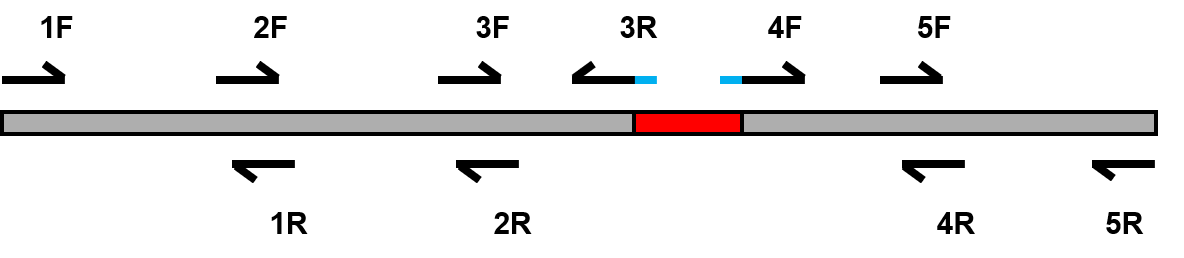

Figure S1: schematic representing T7 phage genome and primer placement for biocontainment. Numbers above primer locations refers to numbers listed in Table S1. Central shaded section (red) represents the omitted capsid gene, *gp10AB*.

Table S2: Strain details used in creating and assaying T7Δcapsid.

| **Strain name** | **Species** | **Source** | **Purpose** |
| --- | --- | --- | --- |
| **DH10β** | *Escherichia coli* | New England Biolabs | Transformation of phage genome for rebooting |
| **TB204/p15A_Capsid** |  | Addgene #230033 | Fluorescently labelled with mCherry; harbouring plasmid allowing replication of T7Δcapsid |
| **TB204/p15A_Control** |  | Addgene #230033 | Fluorescently labelled with mCherry; harbouring plasmid allowing ampicillin resistance without allowing replication of T7Δcapsid |
| **TB205/p15A_Capsid** |  | Addgene #230034 | Fluorescently labelled with GFP; harbouring plasmid allowing replication of T7Δcapsid |
| **TB205/p15A_Control** |  | Addgene #230034 | Fluorescently labelled with GFP; harbouring plasmid allowing ampicillin resistance without allowing replication of T7Δcapsid |

Table S3: Primer details for producing T7Δcapsid. Bolded and capitalised primer sequences indicate inserted homologous overlaps.

| **Primer number** | **5’-3’ binding site (T7 genome)** | **Direction** | **Amplicon length (bp)** | **Sequence** |
| --- | --- | --- | --- | --- |
| **1F** | 1-30 | F | 9999 | tctcacagtgtacggacctaaagttccccc |
| **1R** | 9959-9988 | R |  | tgcagcaataccggaaaggttgtctactgt |
| **2F** | 9970-9999 | F | 10000 | attacgcgatgacagtagacaacctttccg |
| **2R** | 19919-19948 | R |  | acttgtgactccacataggcacatctatga |
| **3F** | 19929-19958 | F | 2988 | atatgtctcctcatagatgtgcctatgtgg |
| **3R** | 22853-22886 | R |  | **GCCCCAAGGGGTTATGCTAG**atttcgaagtctatcagaagttcgaatcgattac |
| **4F** | 24157-24192 | F | 5766 | **CTTCTGATAGACTTCGAAAT**ctagcataac**c**ccttggggcctctaaacgg |
| **4R** | 29879-29908 | R |  | gacatgatggacaagcagggtattgaccct |
| **5F** | 29889-29918 | F | 10059 | gaataacctgagggtcaataccctgcttgt |
| **5R** | 39908-39937 | R |  | agggacacagagagacactcaaggtaacac |

T7Δcapsid NCBI accession number: SAMN57458784
